# Eye-specific voluntary attention can induce a shift of perceptual ocular dominance

**DOI:** 10.1101/2020.08.03.233759

**Authors:** Lili Lyu, Qiu Han, Xin He, Min Bao

## Abstract

Ocular dominance plasticity in adults has been extensively studied in the recent decade. An interocular imbalance of visual input, e.g. monocular deprivation, has been proved to markedly reshape ocular dominance. As visual attention can be eye-specific, dissimilar visual inputs from the two eyes during monocular deprivation inevitably lead attention to be more allocated to the monocular input that conveys relatively intact information. Does the imbalanced attention across the eyes also contribute to reshaping ocular dominance? Here, using a novel “backwards-movie” adaptation paradigm, we showed that prolonged attention to the input in one eye was sufficient to shift perceptual ocular dominance in favor of the unattended eye. Furthermore, the effect was stronger when eye-specific attention was directed to the dominant eye, possibly due to fewer disturbances from the other eye during the adaptation. Taken together, these findings suggest that top-down attention plays an important role in short-term ocular dominance plasticity.

**Statement of Relevance:** An important goal in neuroscience is to understand and take advantage of adaptive neural plasticity. Ocular dominance plasticity resulting from imbalanced visual input signals across the two eyes has been intensively investigated. By developing a novel “backwards-movie” paradigm in which movie images played normally were presented to one eye while movie images of the same episode but played backwards were presented to the other eye, the current study for the first time demonstrates the non-negligible contributions of selective attention in reshaping ocular dominance. These findings expand the homeostatic compensation theory of monocular deprivation by highlighting the contributions of feedback signals. Furthermore, our method could be applied in future work to explore new possibilities in treating adults with amblyopia.

## Introduction

Ocular dominance, which refers to the functional asymmetries of the two eyes, is conventionally thought to be highly plastic in juveniles but hardwired in adults (Wiesel & Hubel, 1963). Recent work on short-term monocular deprivation (MD), however, has demonstrated residual ocular dominance plasticity in the adult visual system by showing that a few hours of MD shifts perceptual and neural ocular dominance in favor of the deprived eye in adults (Bai, Dong, He, & Bao, 2017; Binda et al., 2018; Binda & Lunghi, 2017; Lunghi, Berchicci, Morrone, & Di Russo, 2015; Lunghi, Burr, & Morrone, 2011; Lunghi, Burr, & Morrone, 2013; Lunghi, Emir, Morrone, & Bridge, 2015; Lunghi & Sale, 2015; Ramamurthy & Blaser, 2018; Zhou, Clavagnier, & Hess, 2013; Zhou, Reynaud, & Hess, 2014). This is believed to reflect a homeostatic compensation of the adult visual system.

In typical MD, the non-deprived eye received input of intact images; whereas the deprived eye was presented with either the filtered images in which basic visual properties (such as contrast, phase and spatial frequency) were degraded (Bai et al., 2017; Lunghi et al., 2011; Lunghi et al., 2013; Zhou et al., 2013) or uninformative fractionated images (Ramamurthy & Blaser, 2018). Therefore, it seems that the deprivation of input strength or higher-order visual information is crucial to trigger the shift in ocular dominance. Moreover, neurophysiological evidence has confirmed that short-term MD alters interocular balance in the primary visual cortex, indicating early cortical plasticity (Binda et al., 2018; Lunghi, Berchicci, et al., 2015; Lyu, He, Jiang, Engel, & Bao, 2020; Zhou, Baker, Simard, Saint-Amour, & Hess, 2015).

It is clear that previous literature has demonstrated the key role of the early visual processing in driving the shift of perceptual ocular dominance. However, we noticed that during monocular deprivation, the two monocular pathways received not only dissimilar visual inputs but also distinct attentional modulation. The unfiltered images presented to the non-deprived eye were basically intact natural stimuli thus contained much richer information than those presented to the deprived eye. This allowed the monocular pathway for the non-deprived eye to capture stimulus-driven attention more easily. More than that, goal-driven or top-down endogenous attention was believed to be directed to the non-deprived eye, too. This is because subjects were instructed to engage in everyday activities or watch videos during the deprivation. One question thus arises: Is the imbalanced eye-based attention partially responsible for the shift of ocular dominance? This question cannot be easily addressed using all the existing MD paradigms without any modification, because the effects of imbalanced eye-based attention can be confounded by the effects of imbalanced strength of visual input or higher-order information.

One way to answer this question is to test whether deploying more attention in one eye during the typical short-term MD can modulate the effects of MD. The multiple object tracking (MOT) task is used for this purpose in our pilot experiment (see Augmented-reality experiment in the Supplemental Material available online). The MOT is an attentive tracking task in which subjects keep tracking moving targets for a period of time (Culham, Cavanagh, & Kanwisher, 2001; Zhang, Chen, Yuan, Zhang, & He, 2006), which is often used to continuously control visual attention. In the pilot experiment, we designed a modified version of monocular phase deprivation paradigm (Bai et al., 2017; Lyu et al., 2020) by superimposing the dichoptic MOT stimuli onto the augmented-reality stimuli for MD. Thus during the MD, subjects were presented with equal amount of moving balls in each eye, but tracked only the balls in one of the two eyes (though subjects were not aware that the tracking targets were all in the same eye). If the MOT-induced extra top-down attention in one eye can affect the MD, we would expect the effects of MD to be dependent on which eye the tracking targets were presented to. Unexpectedly, we found that the introduction of the dichoptic MOT abolished the deprivation effect. The finding points to a potential limitation of this type of methods that simply introducing an additional attentive task may inevitably cause the subjects to pay more attention to the task-related stimuli and suppress or ignore the MD background information, making it difficult to test the effects of attention on MD.

To overcome this limitation, we then developed a novel paradigm to directly test whether eye-specific voluntary attention was involved in reshaping ocular dominance. A previous study showed that subjects could continuously attend one episode when two superimposed video episodes were presented in both a binocular and dichoptic mode (Neisser & Becklen, 1975). Based on this finding, we developed a novel “backwards-movie” paradigm in which movie images played normally were presented to one eye while movie images of the same episode but played backwards were presented to the other eye (see Fig. 1a). Since the movie plot of the time-reversed video segment was illogical and hard to follow, selective attention was expected to be predominantly focused on the normally played movie images (presented to the attended eye), allowing us to examine whether prolonged adaptation of top-down, eye-specific attention may lead to changes in ocular dominance. Under the assumption that top-down, voluntary attention enhances the visual processing of the attended eye temporally (Zhang, Jiang, & He, 2012), we hypothesized that the sustained top-down eye-specific attention would boost the response to the signals from the attended eye, resulting in unbalanced response in the two monocular pathways, thus trigger homeostatic mechanism to shift ocular dominance toward the unattended eye. We further examined how the magnitude of the shift in ocular dominance was modulated by different levels of attention. This was realized by manipulating the presence of video sound or its synchronicity with the images.

**Fig. 1.**
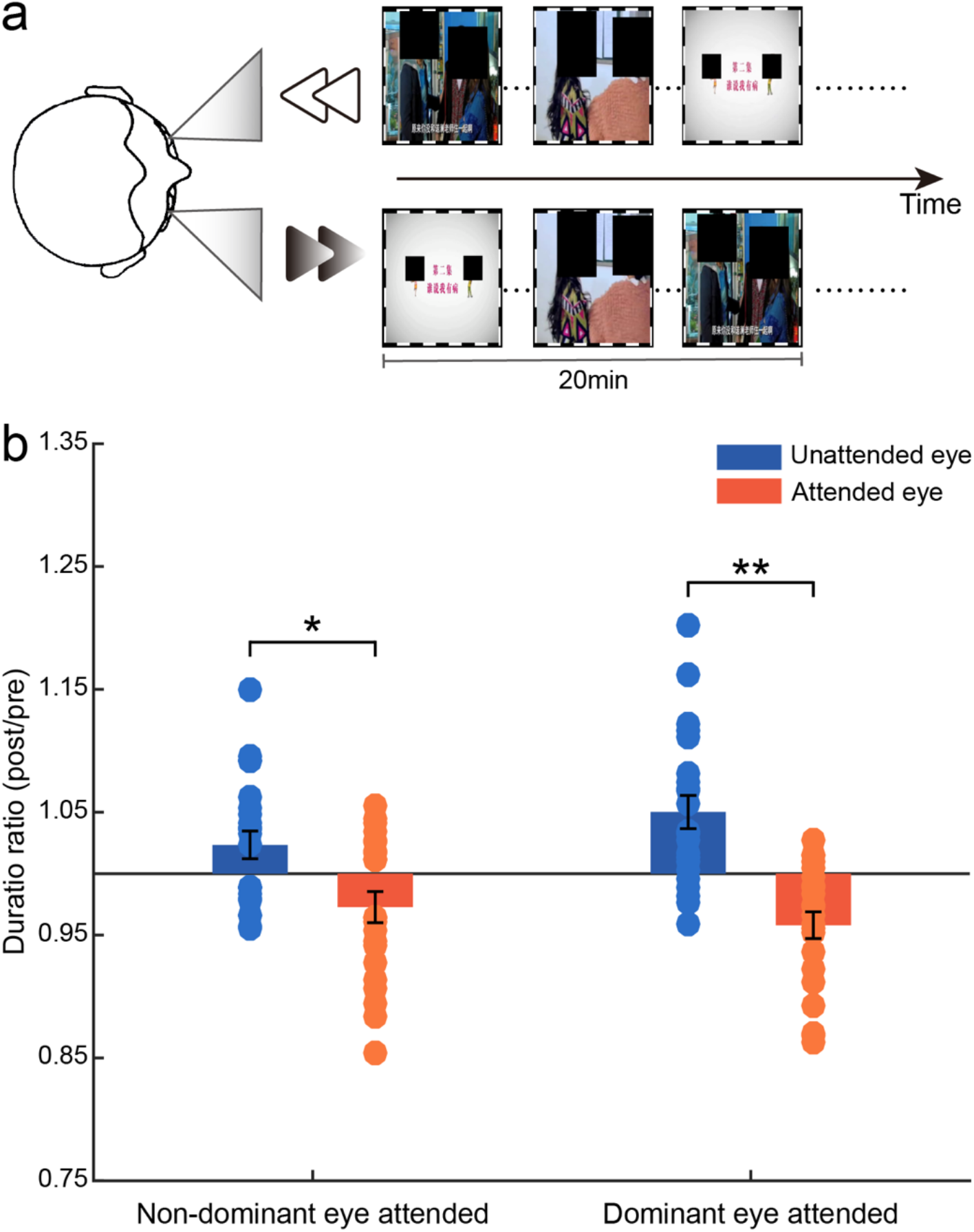
Illustration of the “backwards-movie” paradigm (a) and the mean duration ratios for each eye and each condition for the “Synchrony” condition with 22 subjects (b). Error bars represent standard errors of means. Asterisks represent significant differences between the unattended and attended eye (paired t-test). \**p* < 0.05, \*\**p* < 0.01.

## Methods

### Subjects

Twenty-two young healthy humans (13 females, 9 males; age range 18–25 years) with normal or corrected-to-normal vision participated in the experiment. Six of them only completed one experimental condition (“Synchrony”). All subjects were unaware of the experimental hypotheses and gave informed consent to participate. All experimental procedures were approved by the Institutional Review Board of the Institute of Psychology, Chinese Academy of Sciences and were conducted in accordance with the Code of Ethics of the World Medical Association (Declaration of Helsinki).

### Apparatus

The visual stimuli were programmed in MATLAB and Psychtoolbox (Brainard, 1997; Pelli, 1997). For 8 subjects of the “Synchrony” condition, stimuli were displayed on a gamma-corrected 21-inch Dell P1230 CRT monitor (at the refresh rate of 85 Hz and the mean luminance of 40.5 cd/m^2^). While for other subjects and other conditions, a gamma-corrected 21-inch Sun GDM 5510 CRT monitor (at the refresh rate of 75 Hz and the mean luminance of 42.8 cd/m^2^) was used. The spatial resolution of both monitors was 1024 × 768 pixels. Subjects viewed the stimuli through a mirror stereoscope from a distance of 70 cm, with their heads stabilized in a chinrest. Experiments were conducted in a dark room.

### Stimuli and procedure

#### Binocular rivalry test

Binocular rivalry stimuli were composed of two orthogonal sine-wave grating disks, oriented ±45° (diameter: 1°, spatial frequency: 3 cpd, Michaelson contrast: 80%), presented dichoptically and foveally with a central red fixation point (0.07° in diameter) and a high contrast checkerboard ‘‘frame” (size: 2.5°× 2.5°; 0.25° thick) to facilitate stable binocular fusion.

Each binocular rivalry test was composed of sixteen 60-s rivalry trials. Each trial started with a 5-s blank interval. Then, the rival stimuli were presented for 55 s. Subjects were required to hold down one of the three keys (Left, Right, or Down Arrow) on the keyboard to report their perception (clockwise, counterclockwise, or mixed). The orientation related to each eye was kept constant within a trial, but randomly varied across the trials.

#### Backwards-movie adaptation

During the 1 h of adaptation, subjects passively viewed dichoptically presented movie images, which were surrounded by a high contrast checkerboard ‘‘frame” (size: 11.66° × 18.54°; 0.37° thick). The frame rate of the movie was 25 Hz. The original movie images were presented to one of the two eyes (i.e. the attended eye), while the corresponding backward movie images were presented to the other eye (i.e. the unattended eye). The backward movie images were offline processed by the following steps: dividing the intact movie into a series of 20-min segments; generating a backward copy for each 20-min segment in the MediaEditor software (http://www.aijianji.com/medownload.htm). That is, for each 20-min segment, the backward version of the movie was formally identical to the original one, except for the absence of a logical movie plot.

Our first 6 subjects were only tested for the “Synchrony” condition in which the audio track was always synchronized with the intact movie images. One may argue that the audiovisual integration rather than the eye-specific attention can be responsible for the adaptation effect. To examine the potential contribution of audiovisual integration and manipulate different levels of attention, we added another two different adaptation conditions for the later 16 subjects (we failed to call back the first 6 subjects to test the two additional adaptation conditions). One was the “Asynchrony” condition where the audio track was 5 seconds ahead of the movie image. Studies (Conrey & Pisoni, 2006; Hillock-Dunn & Wallace, 2012) have shown that the time window of audiovisual integration was only several hundred milliseconds wide. Therefore, the 5 seconds interval was far beyond the integration window, making the audiovisual integration very unlikely. The other was the “No-sound” condition where the movie was played without sound.

### Experimental design

This experiment involved two stages: practice and formal experiment.

#### Practice

Before the formal experiment, all subjects practiced three binocular rivalry tests per day (with a 10-min break in between) for 3–7 days. This was to ensure that their performance became stable (Bao, Dong, Liu, Engel, & Jiang, 2018). Previous report (Suzuki & Grabowecky, 2007) has shown that in the first several trials of a day, the measured perceptual eye dominance fluctuated widely. For this reason, in every day of practice, five warm-up rivalry trials were first completed and the data were not analyzed. Perceptual eye dominance was determined by the last three practice sessions with the dominant eye being the one that showed the longer summed phase duration.

#### Formal experiment

The formal experiment had four phases: (1) five warm-up rivalry trials (the data were not analyzed), (2) pre-adaptation binocular rivalry test, (3) 1 h of backwards-movie adaptation and (4) post-adaptation binocular rivalry test.

Each adaptation condition was repeated four times, with each eye attended twice and the sequence counter-balanced. The order of the three conditions (“Synchrony”,”Asynchrony”,”No-sound”) was balanced across subjects.

### Data analysis

To quantify the extend of perceptual dominance of each eye, for the pre- and post-adaptation binocular rivalry tests, we computed the summed phase durations of the exclusively monocular percepts and mixed percepts respectively across all the trials. Then we computed the duration ratio for each eye using the formula (*T_Post_* + *T_Mpost_*/2) / (*T_Pre_* + *T_Mpre_*/2), where *T_Post_* and *T_Pre_* represented the summed predominant durations of one eye during the post- and pre-adaptation tests, and *T_Mpost_* and *T_Mpre_* represented the summed phase durations for mixed percepts during the post- and pre-adaptation tests. If one eye became more dominant in rivalry after adaptation, the ratio would be greater than 1, and vice versa.

## Results

We used a novel “backwards-movie” paradigm to investigate whether top-down, eye-specific attention can induce a shift of ocular dominance. During the adaptation phases of the “Synchrony” condition, the attended eye was always presented with the original movie images that stayed in sync with the movie audio; whereas in the unattended eye, the movie images from a series of 20-min backwards video segments were presented. Since the movie plot of the backwards video segment was illogical and hard to follow, selective attention was expected to be continuously applied to the attended eye. Before and after the 1-h “backwards-movie” adaptation, a binocular rivalry test was used to measure the perceptual eye dominance. For the attended and unattended eye, we respectively divided the posttest predominant duration by the pretest duration to calculate duration ratios.

A2×2 repeated-measures ANOVA with eye dominance (attending dominant or non-dominant eye) and eye status (attended or unattended eye) as factors and paired t-test were used to statistically analyze the results (Fig. 1b). As predicted, the unattended eye showed a higher duration ratio as compared with the attended eye (*F*(1,21) = 19.81, *p* < 0.001, η^2^ = 0.49), indicating increased predominance of the unattended eye after adaptation. And the effect was larger when the dominant eye was attended (*F*(1,21) = 9.03, *p* = 0.007, η^2^ = 0.30). The average duration ratios for the unattended and attended eye were 1.02, 95% confidence interval (CI) = [1.00, 1.05] / 1.05, 95% CI = [1.02, 1.08] and 0.97, 95% CI = [0.95, 1.00] / 0.96, 95% CI = [0.94, 0.98] (see Table 1), which showed quite smaller difference between eyes compared to the values of 1.14 and 0.9 for the deprived and non-deprived eye in Lunghi et al. (2011) study. The small effect found here may be due to the fact that our adaptation duration was only one hour, much shorter than 150 minutes adopted in the previous work (Lunghi et al., 2011).

**Table 1.** Grand average duration ratios (mean [95% CI]) for each condition.

| Condition (number of subjects) | Unattended eye |  | Attended eye |  |
| --- | --- | --- | --- | --- |
|  | Non-dominant | dominant | Non-dominant | dominant |
| Synchrony (22) | 1.02 [1.00, 1.05] | 1.05 [1.02, 1.08] | 0.97 [0.95, 1.00] | 0.96 [0.94, 0.98] |
| Synchrony (16) | 1.03 [1.00, 1.06] | 1.06 [1.03, 1.09] | 0.97 [0.94, 1.00] | 0.95 [0.93, 0.98] |
| Asynchrony (16) | 1.03 [1.00, 1.05] | 1.03 [0.99, 1.07] | 0.97 [0.94, 1.00] | 0.98 [0.95, 1.01] |
| No-sound (16) | 1.01 [0.97, 1.04] | 1.07 [1.03, 1.12] | 1.00 [0.95, 1.04] | 0.94 [0.92, 0.97] |

To disentangle the contribution of audiovisual integration to the observed adaptation effect and manipulate different levels of attention, 16 participants were assigned two additional types of adaptation (“Asynchrony” and “No-sound”, Fig. 2). We performed a three-factor repeated-measures ANOVA: eye dominance (attending dominant or non-dominant eye), eye status (attended or unattended eye) and adaptation type (“Synchrony”,”Asynchrony”,”No-sound”). Table 1 listed the grand average duration ratios for the three conditions. Again, we found a significant main effect of eye status, *F*(1,15) = 25.06, *p* < 0.001, η^2^ = 0.63, and a significant main effect of eye dominance, *F*(1,15) = 10.19, *p* = 0.006, η^2^ = 0.41, but no significant effect of adaptation type, *F(1*.31,19.61) = 1.86, *p* = 0.19, η^2^ = 0.11. Interaction of any two factors was not significant (*p*s > 0.1), but the interaction of three factors was significant (*F*(2,30) = 4.37, *p* = 0.022, η^2^ = 0.23). Simple effect analysis revealed that when the dominant eye was attended, the duration ratio of the deprived eye in the “No-sound” condition was significant larger than that in the “Asynchrony” condition (*p* = 0.048, false discovery rate (FDR) correction), whereas the duration ratio of the non-deprived eye in the “No-sound” and “Synchrony” condition were both smaller than that in the “Asynchrony” condition (*p*s = 0.048, FDR correction). These results indicated that when attending the dominant eye, the magnitude of the shift in ocular dominance for the “No-sound” or “Synchrony” condition was larger than that for the “Asynchrony” condition.

**Fig. 2.**
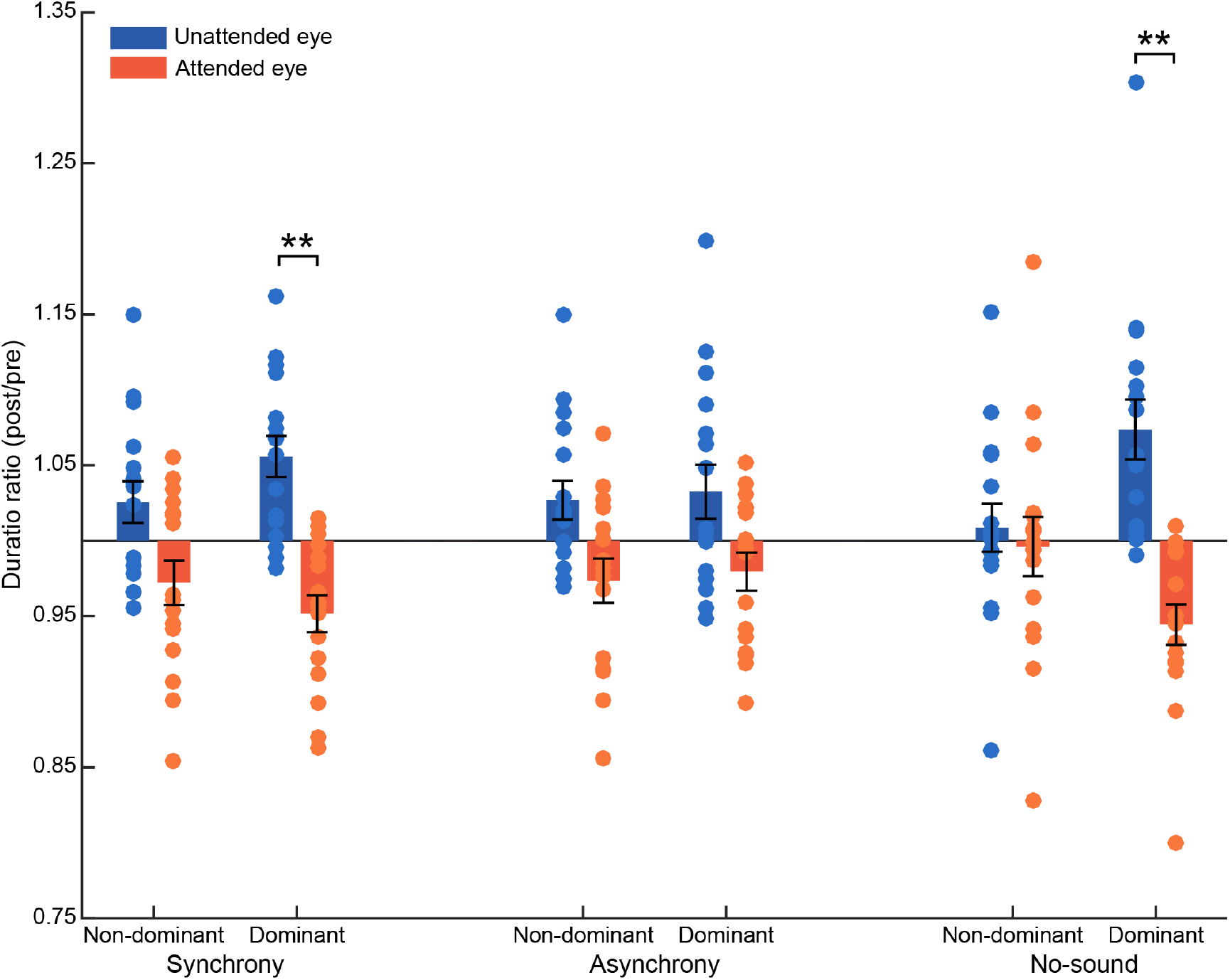
Illustration of the results for the three adaptation conditions. Mean duration ratios for each eye and each condition. Error bars represent standard errors of means. Asterisks represent significant differences between the unattended and attended eye (paired t-test). \*\**p* < 0.01.

## Discussion

Using a novel “backwards-movie” paradigm, we found that 1 h of adaptation of topdown eye-specific attention was sufficient to bias eye dominance in favor of the unattended eye. Because the backwards movie images were identical to the normally played ones except for the reversed frame sequence in each 20-minute segment, both the input energy and higher-order visual information in the two eyes remained generally identical in each segment and so in the entire 1 h of adaptation period. Therefore, neither the low-level input strength nor the higher-order information (e.g. contours) differs across the two eyes. The pattern of the present results supports our hypothesis that the sustained top-down eye-specific attention can boost the response to the inputs of the attended eye during the adaptation period, thus invoke homeostatic plasticity to shift eye dominance toward the unattended eye. For the first time, our findings directly showed the non-negligible contributions of selective attention in producing the effects of monocular deprivation.

Further evidence comes from the results for different adaptation conditions, which showed larger adaptation effects for the “No-sound” or “Synchrony” condition than for the “Asynchrony” condition (when normally played movie was presented to the dominant eye). For the “Synchrony” condition, it was easy for subjects to pay attention to the normally played movie images with the presence of audiovisual integration. And for the “No-sound” condition that lacked auditory cues, subjects had to pay more visual attention to the original images in order to follow the movie plot. Thus, better efficiency or stronger engagement of selective attention could well increase the neural gain for the attended eye in these two conditions. By contrast, under the “Asynchrony” condition, the gain increment was relatively restricted because visual attention applied to the normally played movie images was disturbed by the out-of-sync audio track. As a result, adaptation in the “Asynchrony” condition induced relatively smaller shift of ocular dominance than in the “Synchrony” and “No-sound” conditions.

Additional supporting evidence was that this adaptation effect was larger when the dominant eye rather than the non-dominant eye was attended. When the dominant eye was attended, the normally played movie images were presented to the dominant eye. Thus they could dominate longer than when they were presented to the non-dominant eye. This also meant more chances for the top-down attention to promote the neural gain for the attended eye (rather than the unattended eye) so that a larger effect could be induced. In contrast, when the non-dominant eye was attended, the time-reversed video images were presented to the dominant eye, and dominated longer than when they were presented to the non-dominant eye. In other words, the normally played movie images, to which subjects were eager to pay attention, were now suppressed more often than when the time-reversed video images were presented to the nondominant eye. The consequence was that top-down attention was unable to work in as high efficiency as when the dominant eye was attended, leading to a smaller effect. This result differs from Lunghi et al. (2011) finding that monocular patching induced stronger shifts in eye dominance when the dominant eye was patched (i.e. the nondominant eye was presented with intact images) as compared to the non-dominant eye, suggesting that the underlying mechanisms of adaptation of eye-specific attention and monocular patching were not completely overlapping.

It should be noted that, we did not have any objective measurements to show that subjects did pay more attention to the normally played movie images. According to our pilot experiment, once an additional attentive task was added to the MD background, subjects would pay more attention to the task-related stimuli and suppress or ignore the MD background information. This limitation indicated the difficulty in practice to introduce a behavioral task for measuring the attention level of subjects. Although there were no direct measurements, we believe that some inferences can still be drawn from our experimental design. For each subject, 8 or 12 sessions of the formal experiment were conducted. Different movie episodes from the same series were provided for each session. Attending to the normally played movie images most of the time was helpful for the subjects to follow the episodes, which was also a premise to follow the entire series. Therefore, subjects were motivated to pay more attention to the normally played movie images rather than the time-reversed ones.

Attention has long been shown to modulate the dynamics of binocular rivalry. Cue-mediated involuntary attention can promote the dominance of a rivalry stimulus (Ooi & He, 1999). Secondly, transient top-down eye-specific attention has been shown to prolong dominance durations of an attended target in binocular rivalry (Chong, Tadin, & Blake, 2005) or enhance processing of stimuli presented to the attended eye (Zhang et al., 2012). This eye-specific attentional effect can be reinforced by perceptual training on an attentional cue (Xu, He, & Ooi, 2010) or a feature-based attentional task (Dieter, Melnick, & Tadin, 2016), leading to profound predominance of the attended eye during binocular rivalry. Dieter et al. (2016) showed that the profound predominance of the attended eye was realized both through enhanced eye-specific sensory processing and through strengthened attention to task-relevant features in the attended eye. A novel finding of the present study is that prolonged top-down eye-specific attention can lead to increased predominance of the unattended eye instead of the attended eye. The completely contrary pattern strongly indicates that distinct mechanisms are involved in our adaptation paradigm and eye-based attention training in the previous work.

To speculate how our findings originate at the neuronal level, we still prefer to the framework of homeostatic compensation theory, which was commonly used to explain the findings in recent studies on short-term MD (Binda et al., 2018; Lunghi, Berchicci, et al., 2015; Lunghi et al., 2013; Lunghi, Emir, et al., 2015). Nevertheless, as compared to the existing studies on short-term MD, our results highlight the critical role of attention in reshaping ocular dominance, extending the theory of homeostatic compensation. Attention has been shown to modulate the processing of eye-specific visual information via feedback from higher-level visual areas or frontoparietal cortices (Li, Rankin, Rinzel, Carrasco, & Heeger, 2017; Zhang et al., 2012). Therefore, our results suggest that feedback connections from higher-level areas may play an important role in ocular dominance plasticity, providing new evidence for understanding the underlying mechanism of ocular dominance plasticity in adults. Furthermore, given that our adaptation effect was larger when the dominant eye was attended, it would be intriguing to apply our method in treating adults with amblyopia in future work, as for adult amblyopia the non-amblyopic eye is always the dominant eye.

## Supporting information

see Augmented-reality experiment in the Supplemental Material available online

## Author Contributions

M. Bao conceived the study. All authors contributed to the study design. L. Lyu performed the experiments. L. Lyu and M. Bao analyzed the data and wrote the paper. All authors approved the final version of the manuscript for submission.

## Acknowledgements

This research was supported by the National Natural Science Foundation of China (31871104, 31571112 and 31830037).

