## Supplementary material for "Eye-specific voluntary attention can induce a shift of perceptual ocular dominance": see Augmented-reality experiment in the Supplemental Material available online

### Augmented-reality experiment: Dichoptic MOT during monocular deprivation with pink noise

***Subjects.*** Twelve young healthy humans (5 females, 7 males; age range 20–26 years) with normal or corrected-to-normal vision participated in the experiment. They were naive about the experimental hypotheses and gave informed consent to participate.

***Apparatus.*** The visual stimuli were programmed in MATLAB and Psychtoolbox (Brainard, 1997; Pelli, 1997). For the binocular rivalry measurement, we used a gamma-corrected 27.2-inch LCD monitor (Asus VG278HE) with a resolution of 1920 × 1080 and a refresh rate of 120 Hz to display the stimuli. Subjects viewed the stimuli through a pair of shutter goggles (NVIDIA 3D Vision2 P1431) from a distance of 100 cm, with their heads stabilized in a chinrest. A Photo Research PR-655 spectrophotometer with the sensor attached behind the shutter goggles was used to calibrate the monitor. The mean luminance of the monitor was 20.5 cd/$m^2$ when viewed through the shutter goggles.

For the short-term MD, we adopted two altered reality systems (Bao & Engel, 2019), each of which was comprised of a camera (The Imaging Source) connected to a computer that fed into a head-mounted display (HMD). One system was equipped with a DFK-23UM021 USB3.0 camera (640 × 480 RGB32@60 Hz) connected to a Dell OptiPlex 9010 computer with an NVIDIA Geforce GTX670 graphic processing unit. The other system used the same type of camera connected to a Dell XPS 8700 computer with an NVIDIA GeForce GTX770 graphic processing unit. The HMDs were Sony HMZ-T2 and Sony HMZ-T3 (OLED display, 49.4°in horizontal, 27.8° in vertical, 1280 × 720 pixels).

***Stimuli and procedure.***

*Binocular rivalry test.* The stimuli and procedure were the same as in the backwards-movie experiment.

*MOT during the pink-noise monocular deprivation.* During the adaptation phase, subjects wore the altered reality system and viewed the world through the HMDs. To fit the screen of goggles, we clipped the camera images to a resolution of 640 × 360 pixels and then expanded them to a resolution of 1280 × 720 pixels.

The original camera images were always presented to one of the two eyes (i.e. the non-deprived eye). The camera images presented to the other eye (i.e. the deprived eye) were replaced with pink noises. The power spectra of the pink noises were exactly the same as those of the camera images (their difference was less than $10^{-5}$%). However, the phase spectrum of a pink noise was derived from that of a white noise (randomly selected from 30 pre-defined white noises every 2–5 s). This adaptation condition was referred to as the pink-noise condition (Bai, Dong, He, & Bao, 2017). Camera images were monochrome to speed up the real-time image processing.

Twelve checkboard (square wave grating) balls moved randomly and independently at 4°/s within a virtual square (25° × 25°) on the adapting stimuli background (Fig. S1). They repelled one another and bounced off the edge of the square. A red bull's-eye fixation point was presented foveally and repelled the balls away from the center. For "MOT task" conditions, subjects were required to maintain fixation on this point throughout the tracking period. These balls were dichoptically presented to the two eyes, with six balls in each eye. The non-target balls were 2.3° in diameter, 1.74 cpd in spatial frequency and 50% in Michaelson contrast; whereas the target balls were 3.07° in diameter, 0.89 cpd in spatial frequency and 100% in Michaelson contrast.

The procedure for a typical trial is presented in Fig. S1. Each trial began with the presentation of adapting stimuli alone for 1~4 s, followed by the presentation of a 500Hz sinusoidal tone which was used to inform the subject of the upcoming balls. Then twelve non-target balls appeared and moved in Brownian-like motion. After 1 s the fixation point began to appear and disappear at a frequency of 1 Hz to attract the subject’s attention. The flash lasted for 1 s, then one to three balls turned large (target balls, three trial conditions) for 2 s before reverting to small. Subjects were required to keep tracking the cued balls while maintaining the fixation for the remaining 15 s (tracking period). After the tracking period, a subset of balls turned large for 3 s (probe balls), at which time subjects were required to indicate whether they were identical to the target balls or not by pressing one of the two keys (“Left Arrow” for yes, “Right Arrow” for no). The probe balls were either the same as the target balls, or different by only one ball. The subjects had to make a response within 2 s before the probe balls disappeared. Every incorrect or missed response was accompanied with an auditory feedback (800Hz). For the “Only MOT stimuli with no target” condition, no balls were cued and subjects were instructed to watch the video without paying attention to any balls.

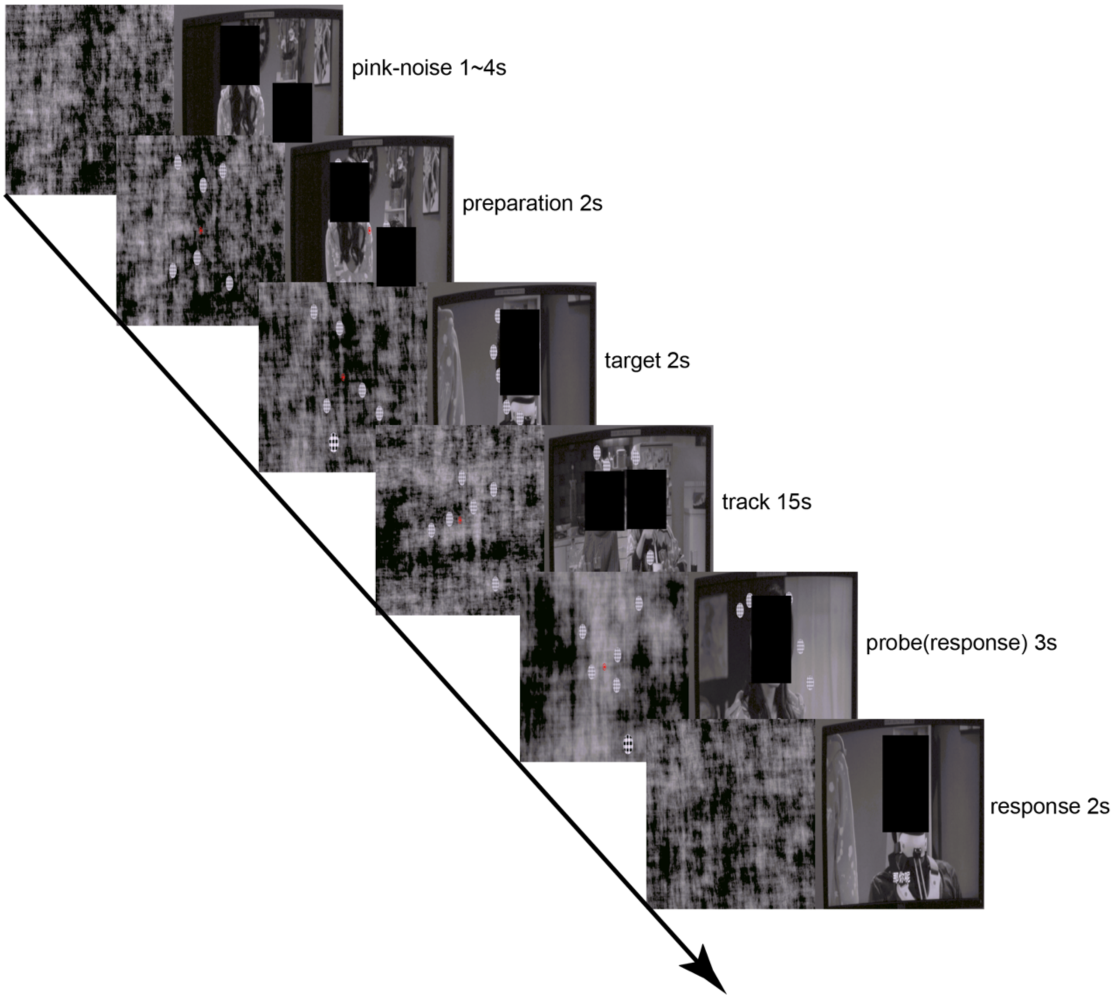


**Fig. S1. Schematic depiction of the MOT task (a one-target trial shown in this example).** (Because of the bioRxiv’s policy, the faces have to be covered, which may affect the image presentation.)

***Experimental design.*** Each subject practiced three binocular-rivalry tests per day for 4 days before the formal experiment. At the same time, subjects also practiced the MOT task on a white noise background. They practiced 60 trials (20 trials for each condition) per day.

In the formal experiment, subjects finished five warm-up rivalry trials first. One binocular rivalry test was measured before and after 120 min of adaptation. There were four adaptation conditions. For the “No MOT stimuli” condition (pure monocular phase deprivation), subjects passively watched the video during the adaptation. For the “Only MOT stimuli with no target” condition, subjects were instructed to watch the video without paying attention to the balls. For the “MOT task with targets on deprived eye” and “MOT task with targets on non-deprived eye” conditions, subjects were instructed to perform the MOT task as accurately as possible while watching the video. The MOT task consisted of 240 trials (80 trials for each condition). There was a 1-min rest every 20 trials, where subjects passively watched the video. After 60 min of adaptation, subjects were instructed to confirm the fusion. It took about 3 h to complete a whole session of experiment. Each subject finished two sessions for each adaptation condition on a separate day, with the sequence for the adaptation condition counter-balanced. The dominant eye was always deprived. The dominant eye was defined as the one that showed the longer summed phase duration in binocular rivalry.

***Data analysis.*** For the binocular rivalry test, the summed phase durations of the exclusively monocular percepts and mixed percepts were respectively calculated across all the trials. The results of two sessions were averaged for each adaptation condition. To quantify the perceptual eye dominance, we calculated an eye ratio index as $(T_{Dep} + T_M/2) / (T_{Ndep} + T_M/2)$, where $T_{Dep}$, $T_{Ndep}$, and $T_M$ represented the summed phase durations for perceiving the stimulus in the deprived eye, the stimulus in the non-deprived eye, and mixed percepts respectively.

A 2 (adaptation condition: no MOT stimuli vs. Only MOT stimuli with no target) ×

2 (test: pre- vs. post-test) repeated measurements ANOVA was performed for the two control conditions. Similarly, for the two "MOT task" conditions, A 2 (adaptation condition: MOT task with targets on deprived eye vs. MOT task with targets on non-deprived eye) × 2 (test: pre- vs. post-test) repeated measurements ANOVA was performed.

***Results.*** Our pilot experiment was based on a dichoptic MOT task against the monocular phase deprivation background. For the two control conditions, the 2 (adaptation condition: no MOT stimuli vs. Only MOT stimuli with no target) × 2 (test: pre- vs. post-test) repeated measurements ANOVA disclosed a significant main effect of test ($F(1,11) = 5.11$, $p = 0.045$, $\eta^2 = 0.32$). No effect of adaptation condition ($F(1,11) = 0.018$, $p = 0.90$, $\eta^2 = 0.002$) or interaction ($F(1,11) = 0.44$, $p = 0.52$, $\eta^2 = 0.039$) was found. In agreement with the previous findings (Bai et al., 2017; Lunghi, Burr, & Morrone, 2011), the monocular phase deprivation induced a significant shift in perceptual eye dominance to the deprived eye (see Fig. S2, no MOT stimuli: $t(11) = 2.25$, $p = 0.046$, $d = 0.65$). When the balls were added to both eyes, the eye ratio index only showed a marginally significant increase after adaptation (see Fig. S2, Only MOT stimuli with no target: $t(11) = 1.97$, $p = 0.074$, $d = 0.57$), suggesting that the introduction of balls might decrease the magnitude of the shift in perceptual eye dominance induced by monocular phase deprivation.

For the two "MOT task" conditions, eye ratio indices were examined in a 2 (adaptation condition: MOT task with targets on deprived eye vs. MOT task with targets on non-deprived eye) × 2 (test: pre- vs. post-test) repeated measurements ANOVA. There was no significant effect of test or adaptation condition, nor an interaction (all *p*s > 0.22). Paired *t*-tests indicated that the eye ratio index showed no change after adaptation for both conditions (all *p*s > 0.15).

To further examine whether the introduction of the ball or the task abolished the deprivation effect, we then performed a 2 × 2 repeated measurements ANOVA for "Only MOT stimuli with no target" and "MOT task with targets on deprived eye" conditions and found a significant main effect of test ($F(1,11) = 11.44$, $p = 0.006$, $\eta^2 =$

0.51). A 2 × 2 repeated measurements ANOVA was also performed for “Only MOT stimuli with no target” and “MOT task with targets on non-deprived eye” conditions and a significant interaction was observed ($F(1,11) = 11.67$, $p = 0.006$, $\eta^2 = 0.52$). The significantly difference between the deprivation effects of these two conditions indicated that the introduction of the tracking task likely abolished the deprivation effect.

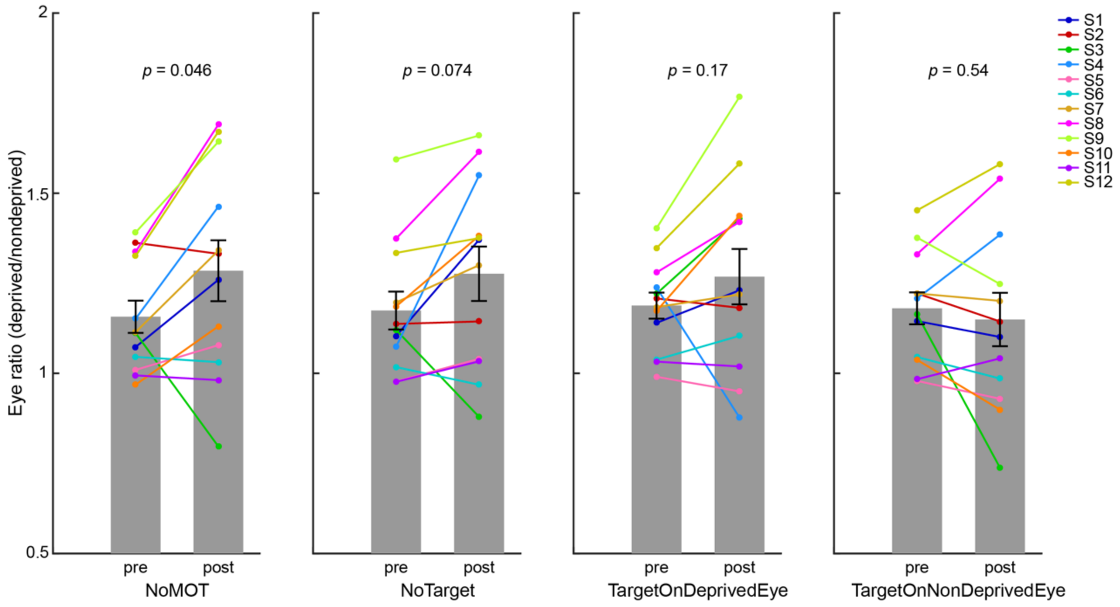


**Fig. S2. Results of the pilot experiment.**

The findings in the pilot experiment support the hypothesis that top-down, eye-specific attention can affect the monocular deprivation effect. MOT is a fairly demanding task. For the two “MOT task” conditions, subjects likely used top-down attention to suppress irrelevant information (i.e. background images) and focus on relevant information (i.e. balls). Since the balls from the two eyes looked the same during the tracking, subjects was unable to efficiently distinguish them (Zhang, Jiang, & He, 2012). As a result, top-down attention was believed to be evenly distributed to the two eyes, leading to weakening or even abolishing the deprivation effect.
